# The mycotoxin Beauvericin exhibits immunostimulatory effects on dendritic cells via activating the TLR4 signaling pathway

**DOI:** 10.1101/2022.01.20.476919

**Authors:** Xiaoli Yang, Shafaqat Ali, Manman Zhao, Lisa Richter, Vanessa Schäfer, Julian Schliehe-Diecks, Marian Frank, Jing Qi, Pia-Katharina Larsen, Jennifer Skerra, Heba Islam, Thorsten Wachtmeister, Christina Alter, Anfei Huang, Sanil Bhatia, Karl Köhrer, Carsten Kirschning, Heike Weighardt, Ulrich Kalinke, Rainer Kalscheuer, Markus Uhrberg, Stefanie Scheu

**Affiliations:** Institute of Medical Microbiology and Hospital Hygiene, Heinrich Heine University Düsseldorf, Düsseldorf, Germany; Institutes of Brain Science, State Key Laboratory of Medical Neurobiology, Fudan University, Shanghai, China; Department of Pediatric Oncology, Hematology and Clinical Immunology, Medical Faculty, Heinrich-Heine University Düsseldorf, Düsseldorf, Germany; Institute of Pharmaceutical Biology and Biotechnology, Heinrich Heine University Düsseldorf, Düsseldorf, Germany; Institute for Transplantation Diagnostics and Cell Therapeutics, Medical Faculty, Heinrich-Heine University Düsseldorf, Düsseldorf, Germany; Institute for Experimental Infection Research, TWINCORE, Centre for Experimental and Clinical Infection Research, a joint venture between the Helmholtz Centre for Infection Research and the Hannover Medical School, Hannover, Germany; Institute of Medical Microbiology, University Hospital of Essen, University of Duisburg-Essen, Essen, Germany; Biological and Medical Research Center (BMFZ), Medical Faculty, Heinrich Heine University Düsseldorf, Düsseldorf, Germany; Institute of Molecular Cardiology, Medical Faculty, Heinrich Heine University Düsseldorf, Düsseldorf, Germany; Julius-Maximilians-Universität of Würzburg (JMU), Würzburg, Germany; Immunology and Environment, Life & Medical Sciences (LIMES) Institute, University of Bonn, Bonn, Germany; Cluster of Excellence - Resolving Infection Susceptibility (RESIST, EXC 2155), Hannover Medical School, Hannover, Germany

## Abstract

Beauvericin (BEA), a mycotoxin of the enniatin family produced by various toxigenic fungi, has been attributed multiple biological activities such as anti-cancer, anti-inflammatory, and anti-microbial functions. However, effects of BEA on dendritic cells remain unknown so far. Here, we identified effects of BEA on murine granulocyte–macrophage colony-stimulating factor (GM-CSF)-cultured bone marrow derived dendritic cells (BMDCs) and the underlying molecular mechanisms. BEA potently activates BMDCs as signified by elevated IL-12 and CD86 expression. Multiplex immunoassays performed on myeloid differentiation primary response 88 (MyD88) and toll/interleukin-1 receptor (TIR) domain containing adaptor inducing interferon beta (TRIF) single or double deficient BMDCs indicate that BEA induces inflammatory cytokine and chemokine production in a MyD88/TRIF dependent manner. Furthermore, we found that BEA was not able to induce IL-12 or IFNβ production in Toll-like receptor 4 (*Tlr4*)-deficient BMDCs, whereas induction of these cytokines was not compromised in *Tlr3/7/9* deficient BMDCs. This suggests that TLR4 might be the functional target of BEA on BMDCs. Consistently, in luciferase reporter assays BEA stimulation significantly promotes NF-κB activation in mTLR4/CD14/MD2 overexpressing but not control HEK-293 cells. RNA-sequencing analyses further confirmed that BEA induces transcriptional changes associated with the TLR4 signaling pathway. Together, these results identify TLR4 as a cellular BEA sensor and define BEA as a potent activator of BMDCs, implying that this compound can be exploited as a promising candidate structure for vaccine adjuvants or cancer immunotherapies.

## Introduction

Dendritic cells (DCs) represent a heterogeneous family of immune cells that link innate and adaptive immunity. They can be classified into two main subtypes: plasmacytoid DCs and conventional DCs, the latter being considered the most potent antigen presenting cells (1-3). DCs are key players in the immune responses, residing in peripheral organs in an immature state and acting as sentinels for a wide array of “danger” signals. These signals include danger-associated molecular patterns (DAMPs) or pathogen-associated microbial patterns (PAMPs), which are recognized by conserved pattern recognition receptors (PRRs) such as the Toll-like receptors (TLRs), RIG-I-like receptors (RLRs) and cytoplasmic DNA receptors (4). Up to date, 10 (TLR1-10) and 12 (TLR1-9, 11-13) functional TLRs are identified in humans and mice, respectively and each is triggered by a distinct set of PAMPs. Based on the distinctive adaptors in these pathways, TLR signaling can be subdivided into two categories: signaling dependent on the MyD88-dependent pathway and signaling dependent on the TRIF-dependent pathway. Most TLRs interact intracellularly with MyD88, except TLR3 which transduces activating signals exclusively via TRIF. Effective TLR4 signaling depends on both adaptor molecules, TRIF and MyD88 (5).

Upon recognition of PAMPs, the TIR domain-containing adaptor proteins MyD88 and/or TRIF are recruited to the TLRs, and initiate signal transduction pathways that culminate in the activation of NF-κB, interferon regulatory factors (IRFs), and mitogen-activated protein kinase (MAP) kinases to upregulate costimulatory molecules (CD40, CD80 and CD86), inflammatory cytokines (e.g. IL-12, IL-6, and TNF), chemokines (RANTES, IP-10, ENA78, etc.) and type I interferons (IFNs) that ultimately protect the host from microbial infection (6, 7). IL-12, composed of p35 and p40 subunits, is the critical factor for Th1 immune responses in the defense against bacterial and viral infection as well as cancer cells (8, 9).

Beauvericin is a cyclic hexadepsipeptide, belonging to the enniatin family and is produced by various fungi, such as *Beaveria bassiana* and *Fusarium spp*. (10, 11). As a mycotoxin, BEA is a very common contaminant of cereal and cereal based products (12, 13), but it is also found in other products such as nuts and coffee (14). Multiple and divers properties of BEA have been reported such as enhancement of pesticide sensitivity (15), anti-bacterial activity against Gram-positive and Gram-negative bacteria (16), anti-viral activity against human immunodeficiency virus type-1 integrase (17), and cytotoxic activity against melanoma cells (18). BEA can also cause cell apoptosis by inducing reactive oxygen species (ROS) production (19). Moreover, BEA shows anti-inflammatory activity in macrophages by inhibiting the NF-κB pathway (20). Although these various properties have been described for BEA, its impact on DCs has not been determined, yet. Here we found that Beauvericin shows immunostimulatory effects on BMDCs via activation of a TLR4 dependent signaling pathway.

## Material and Methods

### Mice

Wild type C57BL/6N and IL-12p40/GFP reporter mice (21) were used for GM-CSF culture of bone marrow cells. OT-II transgenic mice which express an OVA-specific, MHC class II-restricted TCR were used for T cell activation assays. Bone marrow from *Tlr3/7/9*^-/-^ mice and *Tlr4*^-/-^ mice were kindly provided by Prof. Carsten Kirschning. Bone marrow from *Myd88*^-/-^ and *Myd88*^-/-^*Trif*^-/-^ mice was shared by Dr. Heike Weighardt and Prof. Ulrich Kalinke, respectively. No experiments on live animals were performed. Mice were euthanized by cervical dislocation before bone marrow was harvested. The euthanasia method used is in strict accordance with accepted norms of veterinary best practice. Animals were kept under specific pathogen-free conditions in the animal research facilities of the Universities of Düsseldorf, Essen-Duisburg, Bonn and the TWINCORE strictly according to German animal welfare guidelines.

### GM-CSF cell cultures and stimulation conditions

2×10^6^ bone marrow cells were cultured in non-treated 10cm plates (Sarstedt) in 10 ml VLE DMEM (Biochrom) containing 10% heat-inactivated FCS (Sigma-Aldrich), 0.1% 2-mercaptoethanol (Thermo Fischer Scientific), and GM-CSF and kept for 9 days. 10_Jml GM-CSF containing medium was added to the plates at day 3. On day 6 10_Jml medium was carefully removed and centrifuged. The cell pellet was resuspended in 10_Jml medium and added to the dish. On day 9 BMDCs were used for experiments. For cytokine expression analyses, BMDCs were seeded (1×10^6^ cells/well) on a 24-well plate and were stimulated with BEA (purified by the lab of Prof. Rainer Kalscheuer, or purchased from Cayman Chemicals), CpG 2216 (TIB MOLBIOL), LPS (*Escherichia coli* O127:B8, Sigma), cGAMP (InvivoGen), R848 (Alexis Biochemicals), Poly I:C (InvivoGen), or Pam3csk4 (InvivoGen). After 24 hours, cell culture supernatants were collected for cytokine detection.

### Polymyxin B (PMB) neutralization assay

BMDCs were seeded (1×10^6^ cells/ml) on a 24-well plate and were stimulated with BEA (purchased from Cayman Chemicals) or LPS (*Escherichia coli* O127:B8, Sigma) with or without 100µg/ml PMB (InvivoGen). After 16 hours of incubation at 37°C, IL-12p40/GFP expression was analyzed by flow cytometry. Alternatively, after 24 hours, cell culture supernatants were collected and the IL-12p70 and IFNβ content was determined by ELISA assays.

### Flow cytometry and cell sorting

For cell surface staining, fluorochrome-conjugated monoclonal antibodies against mouse CD3e (clone 145-2C11), CD19 (clone 1D3), CD4 (clone RM4-5), and MHC II (clone N5/114.15.2) from BD Biosciences, and CD3e (clone 145-2C11), CD86 (clone GL-1) and CD11c (clone N418) from BioLegend were used. For intracellular staining, cells were first stained for surface markers and then fixed and permeabilized using Intracellular Fixation and Permeabilization Buffer Set (eBioscience) before incubation with fluorochrome-conjugated mAbs against mouse IFNγ (clone XMG1.2, BD Bioscience). Flow cytometry was performed on LSRFortessa (BD Biosciences) or FACSCanto II (BD Biosciences) cytometers. The flow cytometry data was analyzed using FlowJo V10.5. For RNA sequencing experiments live, single, CD3^-^, CD19^-^, CD11c^+^ and MHCII^high^ BMDCs were FACS purified using FACS Aria III (BD Biosciences).

### qRT-PCR

RNA isolation was performed using Macherey-Nagel^™^ NucleoSpin^™^ RNA Plus kit (Macherey-Nagel^™^). Complementary DNA synthesis was done by using the SuperScript^™^ III Reverse Transcriptase (Invitrogen) according to the manufacturer’s instructions. Real-time PCR was performed with 5x MESA Green (Eurogentec) on Bio-Rad CFX 96 Realtime PCR system. Primers used were as follows: β*-actin*: 5’-TGACAGGATGCAGAAGGAGA-3’, 5’-CGCTCAGGAGGAGCAATG-3’; *IL-12p40*: 5’-ACAGCACCAGCTTCTTCATCAG-3’, 5’-TCTTCAAAGGCTTCATCTGCAA-3’; *IFN*β: 5’-CAGGCA ACCTTT AAG CATCA-3’, 5’-CCTTTGACCTTTCAAATGCAG-3’ *TNF:* 5’-TTGAGATCCATGCCGTTG-3’, 5’-CTGTAGCCCACGTCGTAGC-3’; *IL-6:* 5’-CCAGGTAGCTATGGTACTCCAGAA-3’, 5’-GCTACCAAACTGGATATAATCAGGA-3’

### RNA sequencing and analysis

RNA from sorted BMDCs was extracted using the Macherey-Nagel^™^ NucleoSpin^™^ RNA Plus kit (Macherey-Nagel^™^) and quantified using the Qubit RNA HS Assay Kit (Thermo Fisher Scientific). Quality was measured by capillary electrophoresis using the Fragment Analyzer and the ‘Total RNA Standard Sensitivity Assay’ (Agilent Technologies, Inc. Santa Clara, USA). All samples in this study showed high quality RNA Quality Numbers (RQN > 9.4). Library preparation was performed according to the manufacturer’s protocol using the ‘VAHTS^™^ Stranded mRNA-Seq Library Prep Kit’ for Illumina®. Briefly, 500 ng total RNA were used for mRNA capturing, fragmentation, the synthesis of cDNA, adapter ligation and library amplification. Bead purified libraries were normalized and finally sequenced on the NextSeq550 system (Illumina Inc. San Diego, USA) with a read setup of 1×76 bp. The bcl2fastq2 tool was used to convert the bcl files to fastq files.

The reads of all probes were adapter trimmed (Illumina TruSeq) and the clean reads were analyzed using FastQC software to identify potential issues with data quality. The clean reads were then mapped to the mouse reference genome (Mus musculus, GRCm39/mm39) using STAR software. The percentage of uniquely mapped reads were greater than 80%. The uniquely mapped reads to each gene were counted using featureCounts. In order to assess the sample quality, we performed the principal component analysis (PCA) and hierarchical clustering for all samples. No batch effect was detected. The differently expressed genes (DEGs) (|log2FC| >= 1, FDR < 0.05) between non-stimulation and BEA, LPS or BEA with LPS stimulation following the previously described methods (22) were identified using DEseq2 package. DEGs expression was visualized as clustered heat maps using pheatmap package. The functional enrichment analysis (KEGG pathways and GO terms) of DEGs was carried out using enrichR package. Gene Set Enrichment Analysis (GSEA, Version 4.0.3) was used to identify enriched functional gene sets based upon the definitions of the Molecular Signatures Database (23, 24). The included gene set collections were “C2 curated gene sets”, “C5 ontology gene sets” and “C7 immunologic signature gene sets”. An enrichment map of significantly enriched gene sets was produced via Cytoscape (Version 3.8.0) (25) and the GSEA Enrichment Map plugin (26). Since Cytoscape defines a FDR of 0.25 as significant, this value was used as a cut off for inclusion into the network. In the enrichment networks, nodes represent gene sets, while edges represent mutual overlap between gene sets. Genes with overlapping genes and functional annotations were clustered manually to highlight the functional results. These clusters were encircled and labeled with an encompassing terminology. To achieve a simplified and more precise figure all clusters with less than three signatures were discarded from the network.

### Multiplex immunoassay

Cell culture supernatants were assessed for chemokine and cytokine concentrations. The ProcartaPlex Mouse Cytokine & Chemokine Panel 1A 36-plex (Invitrogen by Thermo Fisher Scientific) was used to measure the concentrations of IFNα, IFNγ, IL-12p70, IL-1β, IL-2, TNF, GM-CSF, IL-18, IL-17A, IL-22, IL-23, IL-27, IL-9, IL-15/IL-15R, IL-13, IL-4, IL-5, IL-6, IL-10, Eotaxin (CCL11), IL-28, IL-3, LIF, IL-1α, IL-31, GRO-α (CXCL1), MIP-1α (CCL3), IP-10 (CXCL10), MCP-1 (CCL2), MCP-3 (CCL7), MIP-1β (CCL4), MIP-2 (CXCL2), RANTES (CCL5), G-CSF, M-CSF, and ENA-78 (CXCL5) in cell culture supernatants, according to the manufacturer’s instructions. Plates were read using the Bio-Plex 200 Systems (Bio-Rad, USA).

### ELISA

Cell culture supernatants from BMDCs were analyzed by ELISA for IL-12p70 (R&D) and IFNβ (Invitrogen). Plates were read using a Tecan Sunrise microplate reader at 450 nm, and the background was subtracted at 570 nm.

### Luciferase reporter assay

HEK-293 cells stably expressing TLR4/MD2/CD14 were purchased from Invivogen. Cells were seeded at 3.5 × 10^4^ live cell/well in 96 well plates overnight and were transfected with NF-κB-luciferase reporter plasmid (50 ng/well) and Renilla plasmid (5 ng/well) with transfection reagent jetPRIME (Polyplus-transfection Biotechnology) for 24 hours. Then cells were stimulated with different concentrations of BEA (2.5 μM, 5 μM and 7.5 μM) or LPS (1 μg/ml) as a positive control. After 24 hours of stimulation, the supernatant was discarded, and cells were washed with PBS. 50 µL of lysis buffer (Promega) was added and cells were lysed at room temperature for 15 min on a shaker and luciferase activity was measured with the Dual-Glo Luciferase Assay (Promega) in a Mithras LB 940 multimode microplate reader.

### T cell activation assay

For BMDC / T cell coculture, BMDCs were treated with 2.5 μM, 5 μM, 7.5 μM BEA for 24 hours and then washed twice with PBS to remove residual BEA before use in subsequent culture. Naive CD4^+^ T cells were purified from spleens of OT II mice by MACS (Miltenyi Biotec) according to the manufacturer’s protocol. Briefly, cells were Fc-blocked and incubated with biotinylated anti-CD4 antibodies (BD Pharmingen). Subsequently, magnetic anti-biotin beads (Miltenyi Biotec) were added and CD4^+^ T cells were positively selected by running cells along a MACS magnet. CD4^+^ T cells were labeled with CellTrace Violet (Thermo Fisher Scientific) and afterwards cultured with untreated or BEA treated BMDCs at a 10:1 ratio in 96 well round bottom plates for 3 days and 5 days, respectively. Cell proliferation was measured at day 3 by flow cytometry. At day 5, cell culture supernatants were collected and stored at -80°C. For intracellular detection of IFNγ, Brefeldin A (BD Biosciences) was added to the cells in the last 6 hours before harvesting of the cells at day 5.

### Statistical analysis

GraphPad Prism 9.0 software was used for data analysis. Data are represented as mean ± SEM. For analyzing statistical significance between multiple groups, a one-way ANOVA with Dunnett’s multiple comparisons test was used. For analyzing statistical significance for comparisons of more than two groups with two or more stimulations, two-way ANOVA with Sidak’s multiple comparisons test was used, all p values < 0.05 were considered as statistically significant.

## Results

### BEA activates BMDCs to increase inflammatory cytokine production and costimulatory ligand expression

To study the effects of BEA on BMDCs, cells from IL-12p40/GFP reporter mice were treated with various concentrations of BEA in the presence or absence of suboptimal concentrations of LPS and CpG. BMDC activation was determined by IL-12p40/GFP and CD86 expression. BEA alone potently activated BMDCs leading to enhanced IL-12p40 and CD86 expression (Figure 1A, B). As expected, BEA treatment can also enhance activation of LPS or CpG stimulated BMDCs leading to further increased IL-12p40 and CD86 expression (Figure 1A, B). In addition, we also detected significantly increased production of the inflammatory cytokines IL-12p40, IFNβ, TNF and IL-6 in response to BEA stimulation by Real-time PCR with maximum levels reached at 6 hours post stimulation (Figure 1C). Taken together, BEA can upregulate IL-12 and other pro-inflammatory cytokines together with CD86 levels in BMDCs, indicating that BEA might be a potent BMDC activator.

**Figure 1.**
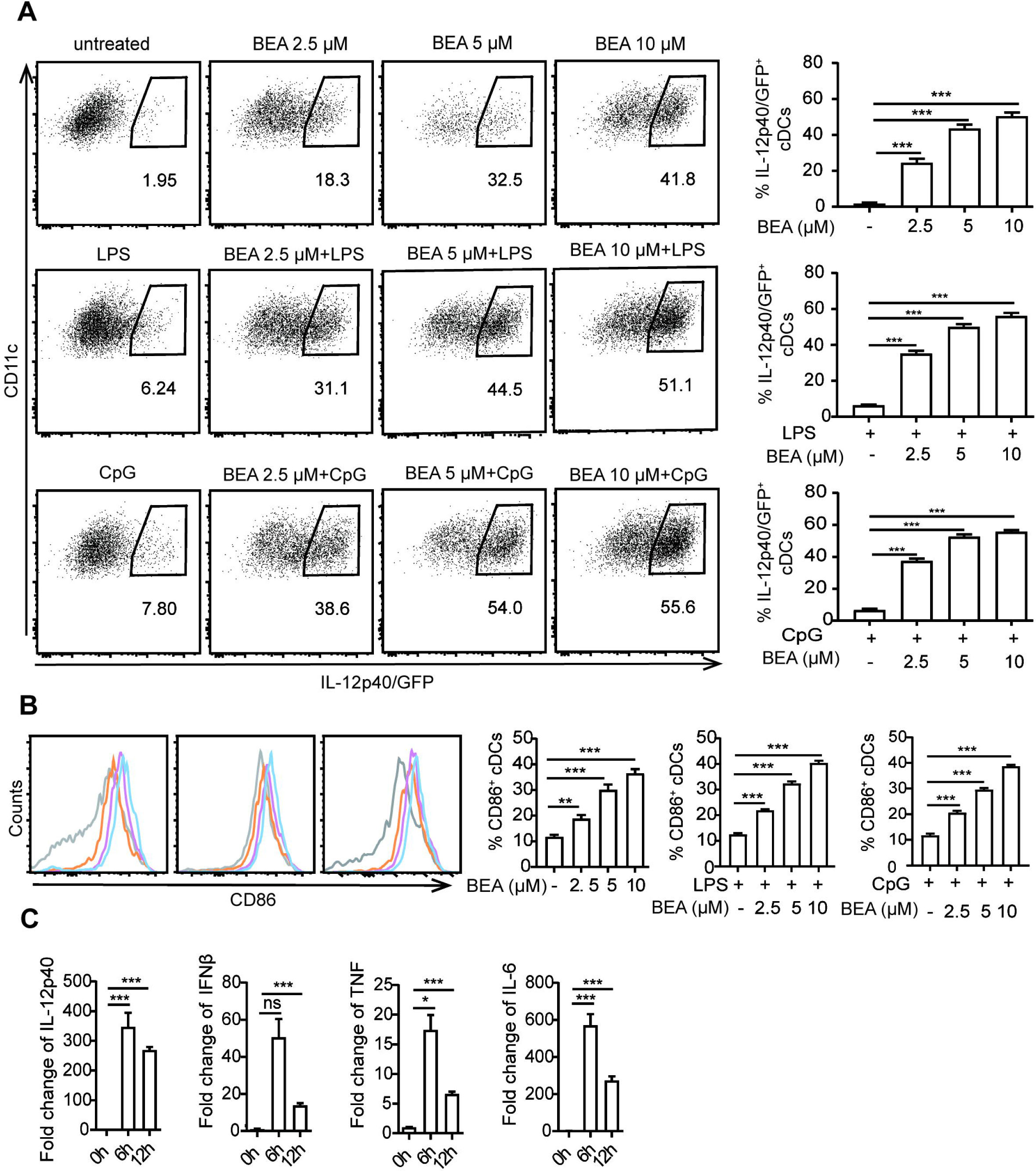
Effects of BEA on BMDCs. **(A, B)** 5×10^5^ BMDCs from IL-12p40/GFP reporter mice were stimulated with indicated concentration of BEA with or without LPS (10 ng/ml) and CpG2216 (0.5 μM) for 16 hours. IL-12p40/GFP **(A)** and CD86 expression **(B)** by BMDCs were detected by flow cytometry. **(C)** 10^6^ BMDCs were stimulated with 5 μM BEA for the indicated time. IL-12p40, IFNβ, TNF and IL-6 were analyzed by Real time PCR. Data are shown as mean ± SEM. Results shown are representative of two to three independent experiments. *p<0.05, **p<0.01, ***p<0.001, ns: not significant.

### BEA promotes DC-mediated CD4^+^ T cell proliferation

Next, we aimed at investigating whether BEA could enhance the ability of BMDCs to induce T cell proliferation. BMDCs were cultured in the presence or absence of various concentrations of BEA for 24 hours and then cells were washed thoroughly as previous studies have shown that BEA significantly inhibits T cell proliferation in TNBS-induced experimental colitis (27). Afterwards, untreated and treated BMDCs were co-cultured with OT II TCR transgenic naive CD4^+^ T cells for 3 days. While untreated BMDCs induced T cell proliferation without BEA stimulation to a certain level, T cell co-culture with BEA treated BMDCs led to increased numbers of T cell divisions (Figure 2A). Furthermore, we also analyzed intracellular IFNγ production of CD4^+^ T cells. While untreated and BEA treated BMDCs showed similar percentages of IFNγ producing CD4+ T cells (Figure 2B), significantly higher IFNγ levels were detected in supernatants of T cells that were co-cultured with BEA activated BMDCs for 5 days than with untreated BMDCs (Figure 2C). The increased amounts of IFNγ in the supernatant of T cells co-cultured with BEA treated BMDCs is due to increased T cell numbers of IFNγ producing T cells but not enhanced capacity for IFNγ production on a per cell basis. This suggests that BEA can enhance the ability of BMDCs to induce T cell proliferation, whereas it does not have an impact on differentiation or induction on cytokine production in individual cells. Taken together, BEA can induce BMDC-mediated CD4+ T cell proliferation.

**Figure 2.**
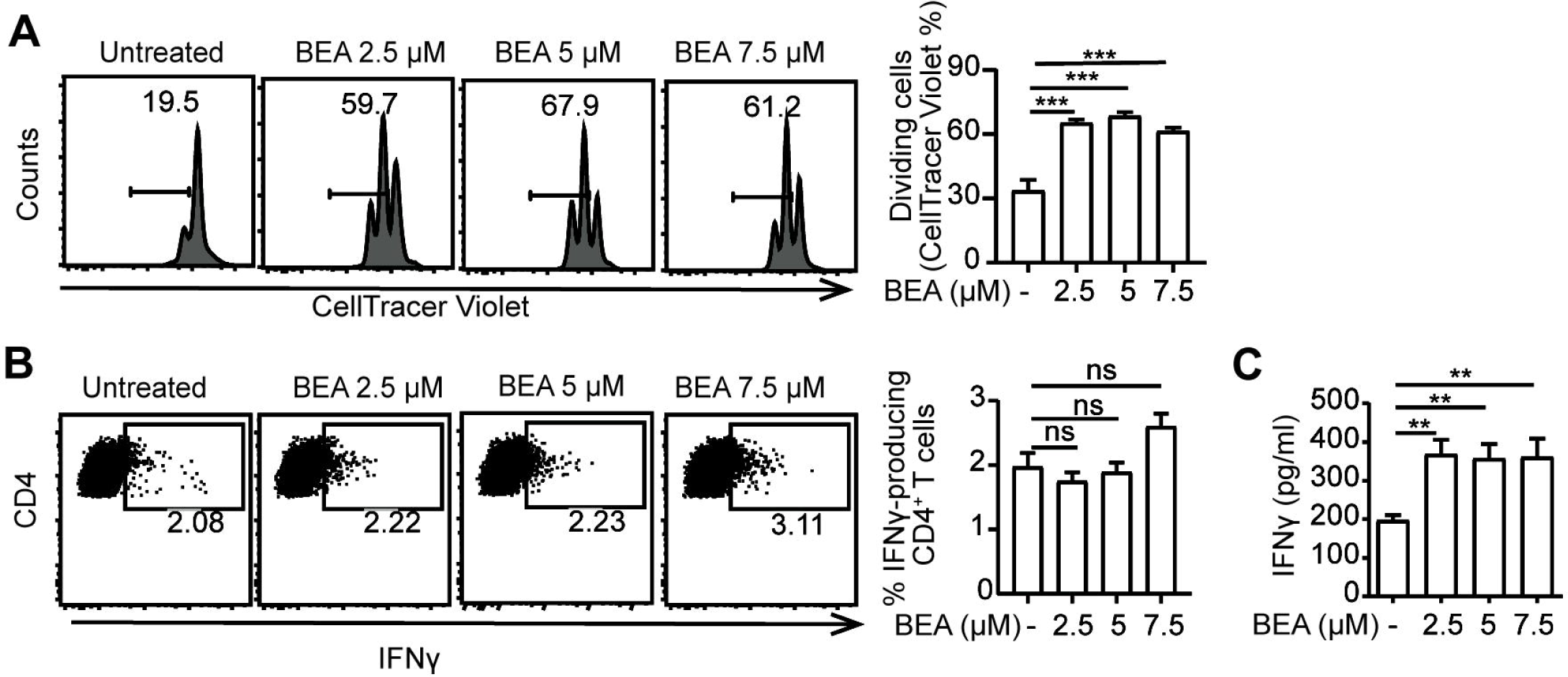
BEA–treated BMDCs enhance T cell proliferation. 10^5^naïve CD4^+^ T cells derived from OT II TCR transgenic mice were labeled with CellTrace Violet and cultured with 10^4^ untreated or BEA-treated BMDCs for 3 days. **(A)** T cell proliferation was analyzed by flow cytometry based on CellTrace Violet dilution. **(B)** 10^5^ naïve OT II CD4^+^ T cells were labeled with CellTrace Violet and cultured with 10^4^ untreated or BEA-treated BMDCs for 5 days. Percentage of IFNγ production by CD4^+^ T cells was measured by intracellular staining. **(C)** IFNγ production in the supernatant was detected by ELISA. Data are shown as mean ± SEM. One representative experiment is shown out of two independent experiments. **p<0.01, ***p<0.001, ns: not significant.

### BEA mediated effects on BMDCs is not due to LPS contamination

The purity of BEA isolated from *Fusarium spp*. by the Institute of Pharmaceutical Biology and Biotechnology (Prof. Rainer Kalscheuer) and BEA purchased from Cayman Chemicals was above 95% as defined by HPLC-UV (data not shown). To further confirm this effect was not a result of endotoxin contamination, BMDCs derived from IL-12p40/GFP reporter mice were stimulated by indicated concentrations of BEA or LPS with or without PMB, which blocks the biological effects of LPS through binding to lipid A (28, 29). After 16 hours of stimulation, IL-12p40/GFP expression by BMDCs was analyzed by flow cytometry. PMB effectively blocked the LPS mediated activation of BMDCs resulting in undetectable IL-12p40 (Figure 3A-B) and IL-12p70 levels (Figure 3C). However, amounts of IL-12p40 and IL-12p70 were comparable after BEA stimulation with or without additional PMB treatment (Figure 3A-C). This result demonstrated that production of IL-12 by BEA treated BMDCs is unlikely to result from any contamination of BEA with LPS.

**Figure 3.**
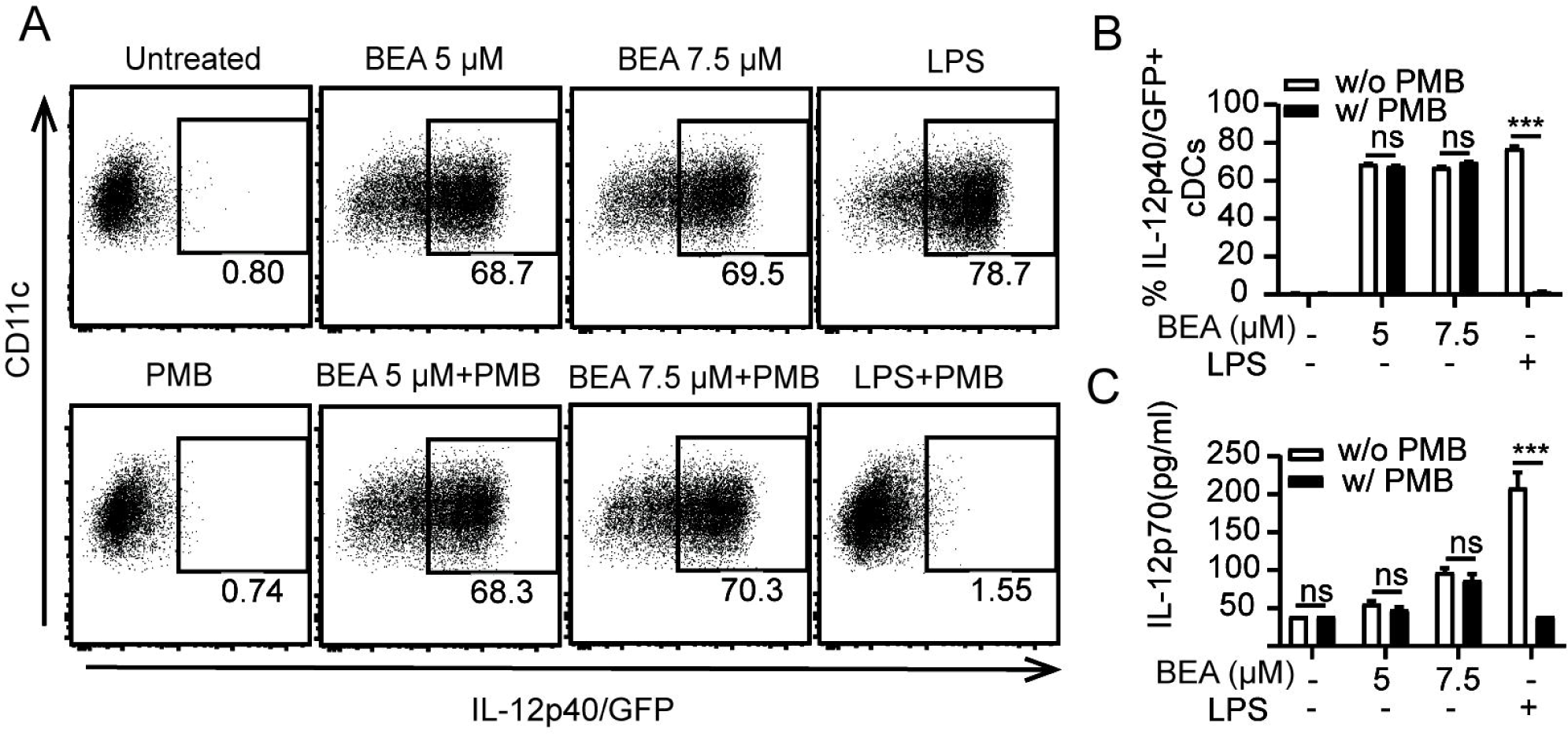
Exclusion of BEA contamination with LPS. **(A, B)** 5×10^5^ BMDCs derived from IL-12p40/GFP reporter mice were stimulated with 5 µM BEA or LPS (10 ng/ml) with or without PMB (100 ng/ml). After 16 hours of stimulation, IL-12p40/GFP and CD86 expression of BMDCs were analyzed by flow cytometry. **(C)** 10^6^ BMDCs derived from IL-12p40/GFP reporter mice were stimulated by 5 µM BEA or LPS (10 ng/ml) with or without PMB (100 ng/ml). After 24 hours of stimulation, supernatants were analyzed for IL-12p70 by ELISA. Results shown are representative of two independent experiments. ***p<0.001, ns: not significant.

### BEA induces BMDC cytokine production in a MyD88 and TRIF dependent way

MyD88 and TRIF are critical adaptors for TLR induced production of pro-inflammatory cytokines such as IL-12, TNF, IL-6 and IFNβ by DCs (30, 31). Therefore, we aimed to investigate whether IL-12 and IFNβ production by BEA stimulation are MyD88 or TRIF-dependent. To this end, BMDCs were generated from *Myd88*^-/-^ and *Myd88*^-/-^ *Trif*^-/-^ mice and stimulated with BEA, LPS or cGAMP, the latter serving as a positive, MyD88/TRIF-independent stimulation control. After 24 hours stimulation, cell supernatant was collected to assess IL-12 and IFNβ production by ELISA. In this experiment, cGAMP can induce IFNβ production in WT and *Myd88*^-/-^ or *Myd88*^-/-^ *Trif*^-/-^ BMDCs. As expected, neither *Myd88*^-/-^ nor *Myd88*^-/-^ *Trif*^-/-^ BMDCs released detectable amounts of IL-12 upon LPS stimulation. Production of IFNβ was significantly diminished but still detectable in *Myd88*^-/-^ BMDCs, whereas it was undetectable in *Myd88*^-/-^ *Trif*^-/-^ BMDCs. Similarly, BEA did not induce IL-12p70 production in either *Myd88*^-/-^ BMDCs or *Myd88*^-/-^ *Trif*^-/-^ BMDCs while IFNβ production was significantly decreased in *Myd88*^-/-^ BMDCs and undetectable in *Myd88*^-/-^ *Trif*^-/-^ BMDCs (Figure 4A). Furthermore, we determined production of other cytokines and chemokines by multiplex immunoassay. Production of the inflammatory cytokines TNF, IL-6, IL-1β, IL-18, IL-27 and IL-10 (Figure 4B) and the chemokines GRO-α, MCP-3, ENA-78, MIP-1β and RANTES (Figure 4C) was significantly decreased in BEA simulated *Myd88*^-/-^ BMDCs and even lower in BEA simulated *Myd88*^-/-^ *Trif*^-/-^ BMDCs. However, production of IP-10 induced by LPS in *Myd88*^-/-^ BMDCs was similar to WT BMDCs, but was markedly decreased in LPS stimulated *Myd88*^-/-^ *Trif*^-/-^ BMDCs. Such findings are consistent with studies reporting that expression of IP-10 by LPS-stimulated bone-marrow-derived macrophages is mediated through a TRIF-dependent but MyD88-independent pathway (32). Interestingly, similar results were observed in BEA stimulated *Myd88*^-/-^ BMDCs and *Myd88*^-/-^ *Trif*^-/-^ BMDCs. Thus, the effects of BEA on BMDCs cytokine and chemokine expression profiles are mediated via activation of MyD88 and TRIF signaling pathways as similarly detected after LPS-stimulation.

**Figure 4.**
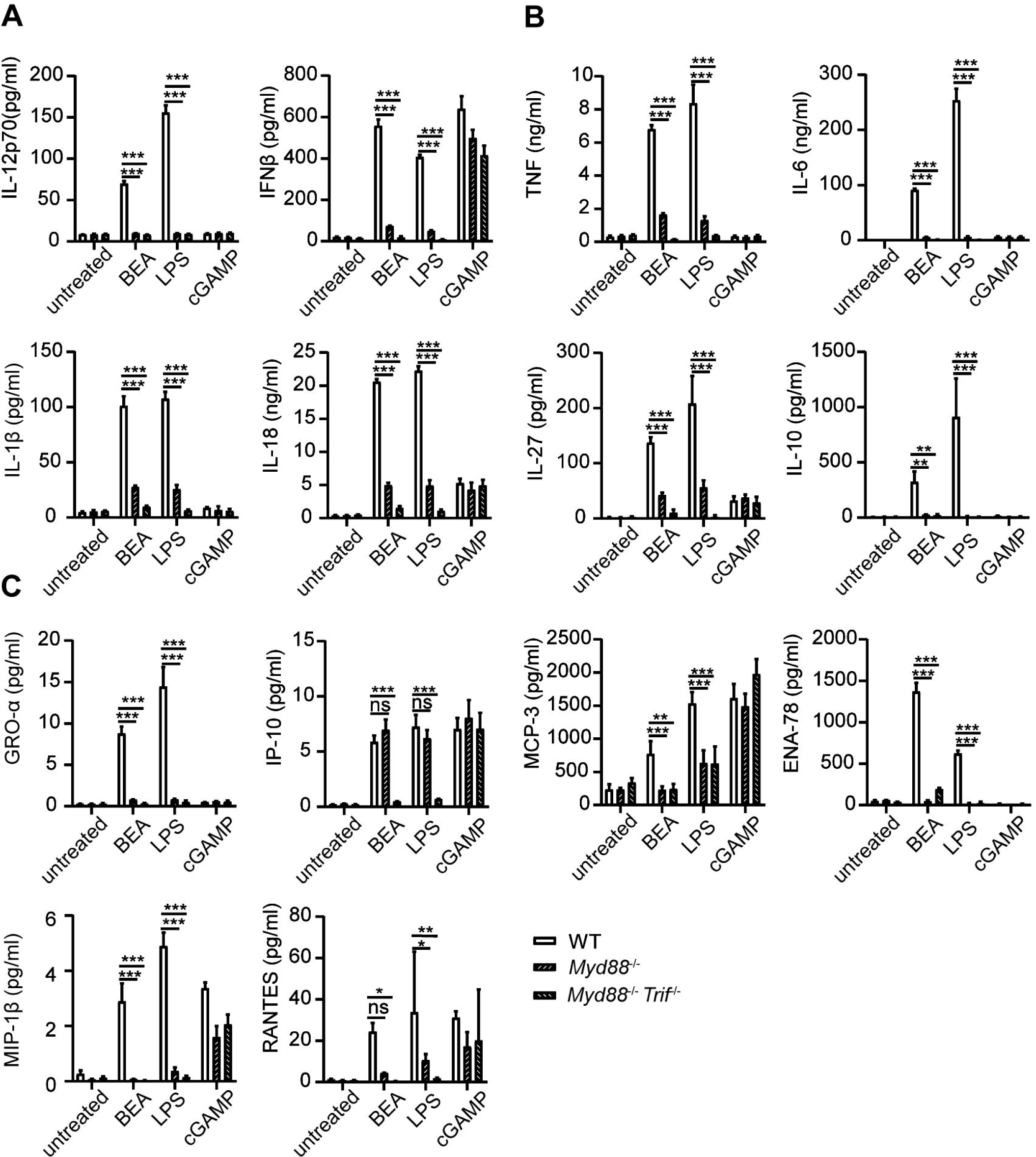
BEA promotes BMDC activation in a MyD88/TRIF dependent manner. 10^6^BMDCs from WT, *Myd88*^*-/-*^ and *Myd88*^-/-^ *Trif*^-/-^ mice were stimulated with 5 µM BEA, LPS (10 ng/ml) and cGAMP (10 ng/ml) as control. After 24 hours of stimulation, supernatants were analyzed for IL-12p70 and IFNβ production by ELISA **(A)**, and inflammatory cytokine (TNF, IL-6, IL-1β, IL-27, IL-10) **(B)** and chemokine production (GRO-alpha, IP-10, MCP-3, ENA-78, MIP-1α and RANTES) **(C)** by Multiplex immunoassays. Results shown are representative of two independent experiments. *p<0.05, **p<0.01, ***p<0.001, ns: not significant.

### BEA activates BMDCs in a TLR4-dependent way

It has been shown that the TLR4 signaling pathway not only depends on the presence of the MyD88 signal adaptor protein but also the TRIF signal adaptor protein (31). As we observed that both, MyD88 and TRIF are involved in BEA induced BMDC cytokine production, we next sought to determine whether BEA activates BMDCs in a TLR4 dependent manner. To this end, we stimulated WT and *Tlr4*-deficient BMDCs with BEA in the presence or absence of CpG or LPS for 24 hours. Measurement of IL-12p70 and IFNβ in the supernatant by ELISA showed that LPS and BEA did not induce IL-12p70 and IFNβ production in *Tlr4*-deficient BMDCs (Figure 5A, B). In contrast, CpG induced similar amounts of IL-12p70 and IFNβ in both WT and *Tlr4*-deficient BMDCs while BEA co-treatment with CpG failed to induce more cytokine production as compared to *Tlr4*-deficient BMDCs stimulated with CpG alone, suggesting these effects of BEA on BMDCs are TLR4 signaling dependent.

**Fig 5.**
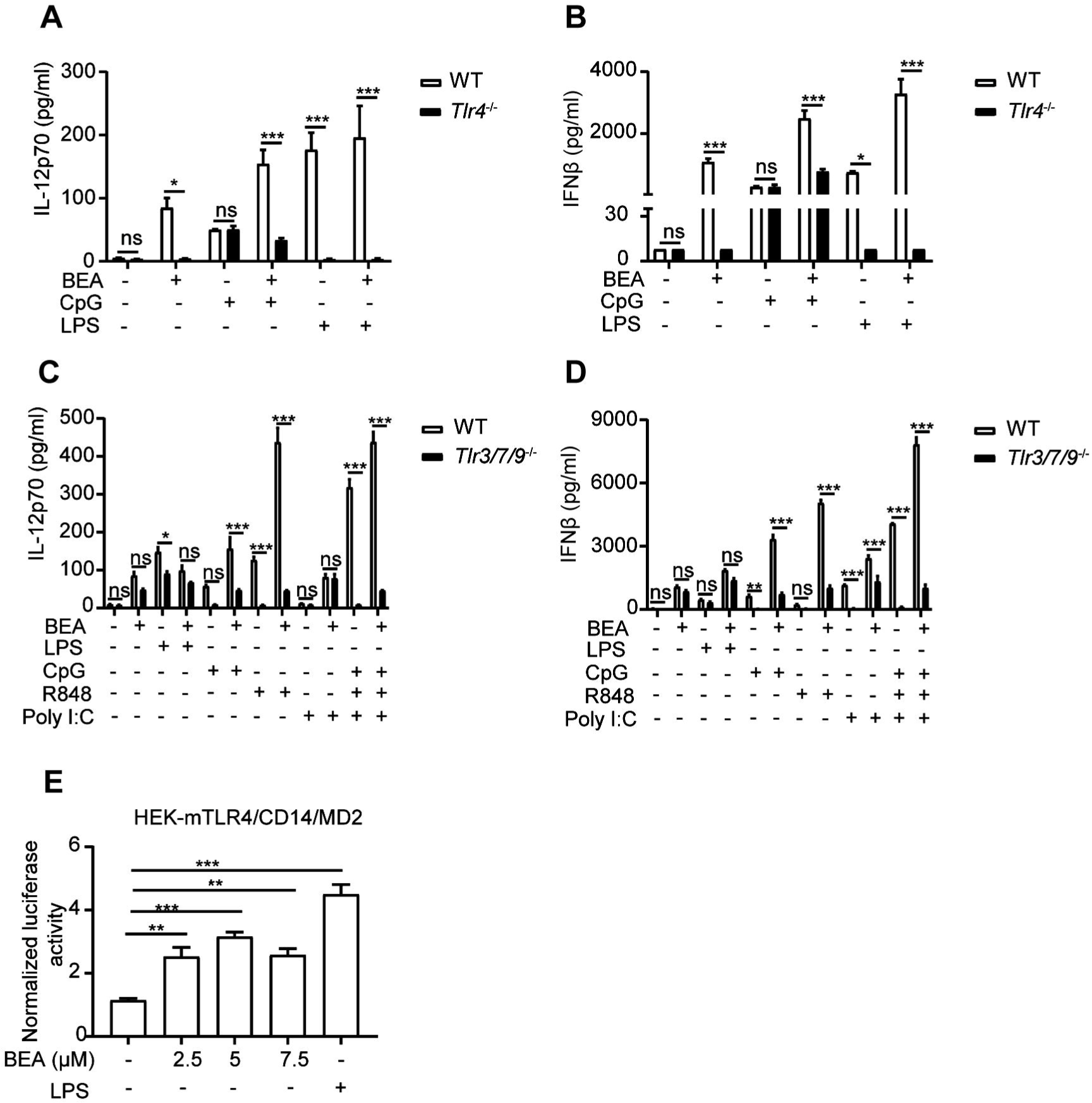
BEA activates BMDCs via TLR4. **(A, B)** 10^6^ BMDCs derived from WT and *Tlr4*^-/-^ mice were stimulated with 5 µM BEA with or without LPS (10 ng/ml) and CpG (0.5 μM). After 24 hours of stimulation, supernatants were analyzed for IL-12p70 and IFNβ by ELISA. **(C, D)** 1×10^6^ BMDCs derived from WT and *Tlr3/7/9*^-/-^ mice were stimulated with 5 µM BEA with or without LPS (10 ng/ml), CpG (0.5 μM), R848 (1 μg/ml), or Poly I:C (25 ng/ml). After 24 hours of stimulation, supernatants were analyzed for IL-12p70 and IFNβ by ELISA. Results shown are representative of two independent experiments. **(E)** 3.5×10^4^ HEK-293 cells stably expressing mTLR4/CD14/MD2 were transiently transfected with firefly luciferase NF-κB reporter and Renilla plasmids. After 24 hours, transfected HEK-293 were treated with indicated concentrations of BEA and LPS (1 μg/ml) as positive control and induction of NF-κB was determined by luciferase activity. Results shown are representative of three independent experiments. *p<0.05, **p<0.01, ***p<0.001, ns: not significant.

Furthermore, to investigate whether BEA can activate other TLR signaling pathways, WT and BMDCs with a triple deficiency of TLR3, 7 and 9 were stimulated by BEA with or without CpG (TLR9), R848 (TLR7) or Poly I:C (TLR3). Consistent with current knowledge, CpG and R848 can significantly induce IL-12p70 and IFNβ production in WT, but not in *Tlr3/7/9*-deficient BMDCs. In addition, we found that Poly I:C did not induce IL-12p70 production, which is consistent with previous reports (30). In contrast, BEA induced similar amounts of IL-12p70 and IFNβ in WT and *Tlr3/7/9* deficient BMDCs (Figure 5 C, D).

To further determine if BEA could directly activate TLR4-mediated signaling, we stimulated HEK-293 cells stably expressing mTLR4/CD14/MD2 and transiently expressing the NF-κB-luciferase reporter and Renilla gene with various concentrations of BEA or LPS as a positive control and measured NF-κB activation. LPS treatment significantly induced NF-κB activation, which was similarly observed after BEA treatment (Figure 5E). Taken together, our data indicate that BEA activates BMDCs via a TLR4 dependent signaling pathway.

### BEA induces transcriptional changes associated with TLR signaling and chemokine signaling pathways

To define the underlying mechanisms by which BEA activates BMDCs, we used whole-genome RNA sequencing (RNA-seq) to detect genome wide differences in gene expression of BMDCs treated with or without BEA in an explorative study. MHC II^high^ CD11c^+^ BMDCs were sorted by flow cytometry followed by stimulation with BEA or LPS alone or BEA combined with LPS for 4 hours. Control samples were left untreated. PCA revealed that the four treatment groups cluster separately and that combined BEA with LPS treatment clusters in close proximity to that of BEA stimulation alone (Figure 6A). Similarly, heatmap and hierarchical clustering show that gene expression induced by BEA is different from LPS stimulation. Combined BEA and LPS stimulation induces a similar differential gene expression as BEA stimulation sharing differential regulation of endolysosome related gene expression (Lamp1, Lamp2, Lamtor3, CSTB, Vps35 and Mcoln1), cellular metabolism gene expression (HK3 and Fasn), mitochondrial gene expression (Polrmt, Slc25a29), autophagy gene expression (rptor) and transcription regulation (Zfp446, H4c3 and foxf2) (Figure 6B). KEGG pathway and GO analyses for BEA treated versus untreated BMDCs were enriched in those involved in “the innate TLR pathway”, “the MyD88 mediated pathway”, “the cytokine signaling pathway”, “the chemokine signaling pathway”, “response to lipopolysaccharide” and “regulation of interleukin-6 production”, amongst others, which further confirmed our Multiplex results (Figure 6C). Using Cytoscape to visualize molecular interaction networks we could show that BEA, LPS and BEA together with LPS similarly induced regulation of clusters related to DCs activation and cell cycle progression (Figure 6D and Supplemental Figure 1). In contrast, BEA led to additional clusters associated with cellular metabolism, T-cell activation, complement activation, type I IFN response, vaccine response, JAK-STAT signaling, ribosomes, translation, and autophagy/receptor recycling (Figure 6D). Moreover, BEA with LPS synergistically and additionally induced B cell antibody production networks and innate immune response (Supplemental Figure 1). Taken together, our results indicate that BEA activates BMDCs via a TLR4 dependent signaling pathway, but induces a gene expression profile different from LPS.

**Figure 6.**
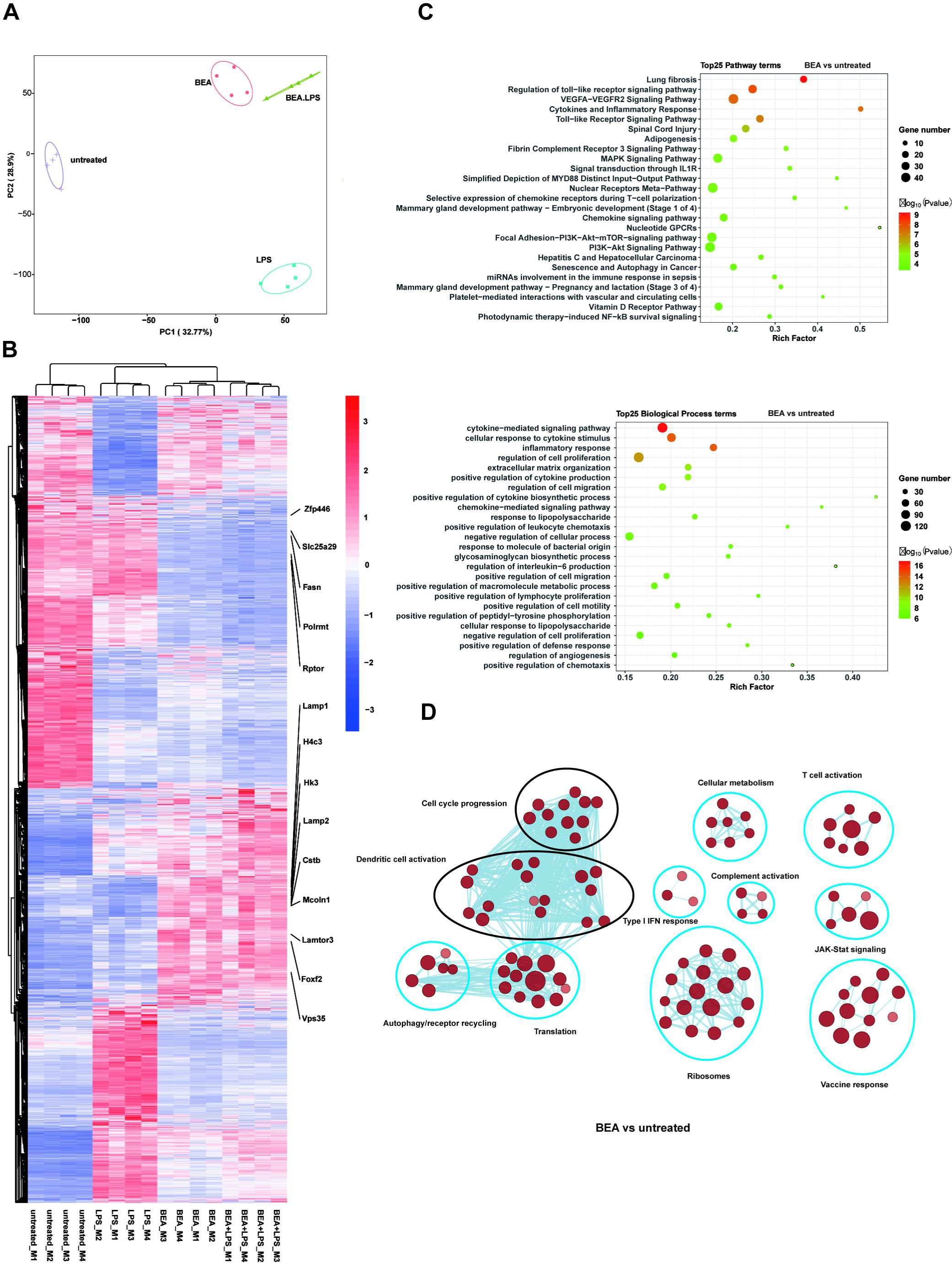
BEA promotes transcriptional changes associated with chemokine and cytokine production and TLR signaling pathway activation but are distinct from LPS stimulation. MHC II^high^ CD11c^+^ BMDCs sorted by flow cytometry were followed by stimulation with BEA (5 µM) or LPS (1 μg/ml) alone or BEA combined with LPS for 4 hours. **(A)** PCA of the quadruplicate biological replicates of each condition. **(B)** Heatmap showing expression profile for 4,015 genes that were found to be significantly regulated in at least one of the comparisons using untreated as baseline condition. **(C)** Enriched Reactome pathways (upper plot) and biological process (lower plot) in DEGs in BEA treated BMDCs compared with untreated BMDCs. **(D)** Cytoscape representation of significantly enriched signatures in BEA treated BMDCs compared with untreated BMDCs.

## Discussion

BEA is a natural product found in various toxigenic fungi, for which several biological effects have been reported, such as cytotoxic, apoptotic, anti-cancer, anti-microbial, insecticidal, and nematicidal activities (14). Moreover, BEA has been reported to exhibit anti-inflammatory activity in macrophages by inhibiting the NF-κB pathway and in an experimental colitis model by inhibiting activated T cells (20, 27). However, little is known about the effect of BEA on DCs. In this study, we showed for the first time that BEA activates GM-CSF-cultured BMDCs, inducing inflammatory cytokines such as IL-12, IFNβ, TNF, IL-6 together with CD86 expression in a MyD88 and TRIF-dependent way. Furthermore, BEA can enhance the ability of BMDCs to induce T cell proliferation, whereas it does not have an impact on differentiation or induction on cytokine production in individual cells. The purity of isolated and commercial BEA is above 95% and our PMB-blocking experiments also exclude any possibility of endotoxin contamination.

TLRs are crucial activating receptors on antigen presenting cells including macrophages and DCs. Upon recognition of PAMPs or DAMPs, they can induce a variety of cellular responses including production of inflammatory cytokines, chemokines, and type I IFNs. TLR signaling consists of at least two distinct pathways: a MyD88-dependent pathway that leads to the production of inflammatory cytokines, and a MyD88-independent pathway associated with the induction of IFNβ (5, 33, 34). Signaling downstream of most of TLRs is MyD88-dependent, except for signaling downstream of TLR3, which is exclusively TRIF-dependent. TLR4 signals through both, the MyD88- and TRIF-dependent pathway to induce inflammatory cytokines, chemokines and type I IFNs production (30). To explore the mechanism by which BEA activates BMDCs, we analyzed inflammatory cytokine and chemokine production by BMDCs derived from *Myd88*^-/-^ or *Myd88*^-/-^ *Trif*^-/-^ mice after BEA stimulation. Production of cytokines and chemokines in response to BEA stimulation was strongly diminished in *Myd88*^-/-^ BMDCs and almost undetectable in *Myd88*^-/-^ *Trif*^-/-^ BMDCs. Thus, these results suggest BEA activates BMDCs using signaling pathways that are both MyD88- and TRIF-dependent. Thus, we reasoned that BEA activates BMDCs via activating the TLR4 signaling pathway. To test this, we analyzed the release of cytokines from *Tlr4*^-/-^ BMDCs. BEA significantly decreased IL-12p70 and IFNβ production by *Tlr4*^-/-^ BMDCs. Consistently, Luciferase Reporter Assay shows that BEA significantly induced NF-κB activation in HEK-293 cells stably expressing TLR4/MD2/CD14. Moreover, RNA sequencing and GO analyses showed that BEA-treated BMDCs activate pathways related to TLR signaling, cytokines and inflammatory response, chemokine signaling, and IL-10 anti-inflammatory signaling, which were similarly activated in LPS-treated BMDCs. However, also marked differences exist between BEA-treated BMDCs and LPS-treated BMDCs. BEA-treated BMDCs show regulation of various signatures associated with cellular metabolism, T cell activation, complement activation, type I IFN response, vaccine response, JAK-STAT signaling, ribosomes, translation, and autophagy/receptor recycling, which was not found to the same extend in LPS-stimulated BMDCs. These differences could be attributed to the different affinity of TLR4 to BEA and LPS or by additional molecular targets of BEA within the cells. Of note, heat-killed conidia of *Aspergillus fumigatus* have been reported to activate TLR4 signaling to induce inflammatory cytokine production (35). However, which component of this fungi is responsible for TLR4 activation was not elucidated. It is tempting to speculate that BEA or a derivative thereof produced by this fungus is responsible for this TLR4 stimulating activity, but this remains to be elucidated in future studies.

It has been reported that BEA shows cytotoxicity on human DCs derived from human umbilical cord blood CD14+ monocytes. Furthermore, BEA can affect LPS-induced DCs maturation by decreasing CCR7 expression and increasing IL-10 production (36), whereas, effects of BEA alone on human dendritic cell activation remain unknown. To determine whether BEA can activate human DCs is the aim of future studies. Furthermore, we found BEA pre-treated BMDCs could enhance T-cell proliferation, whereas no difference of T-cell proliferation was observed when BEA was present in the co-culture of BMDCs together with T cells (data not shown). This could be caused by direct inhibition of T-cell proliferation by BEA resulting in a neutralization of BMDC-mediated T-cell proliferation (27). Of course, further studies need to be done to verify effects of BEA *in vivo*. In addition, BEA has been reported to exhibit anti-inflammatory activity in macrophages by inhibiting the NF-κB pathway (20). To determine why DCs and macrophages react differently to BEA is another task of future studies. Also, further studies need to be done to define the molecular mode of action of BEA-mediated TLR4 stimulation. Direct binding of BEA to TLR4 needs to be tested and if applicable the TLR4 domains involved need to be identified. Alternatively, BEA-mediated TLR4 signalling could be activated via the release or induction of endogenous proteins serving as ligands for TLR4 such as Mrp8 (37), heat shock proteins (HSP60, 70, Gp96) (38) and high mobility group box 1 protein (HMGB1) (39).

Adjuvants are defined as molecules or formulations that enhance the efficacy of vaccines without directly participating in the protective immunity. In recent decades, a variety of preclinical and clinical studies have shown that purified TLR agonists could be exploited as adjuvants to enhance adaptive responses during vaccination (40, 41). Monophosphoryl lipid A (MPLA), a TLR4 agonist purified from *Salmonella minnesota* LPS has been used as adjuvant in different vaccines against human papillomavirus (HPV) and hepatitis B virus (HBV) infections (42). Moreover, MPLA is the only TLR4 agonist that has been clinically tested as an adjuvant for cancer vaccines (43, 44). In our study, BEA potently activated DCs inducing a range of inflammatory cytokines and chemokines in addition to MHC II upregulation. By means of a cell directed delivery of BEA a specific activation of DCs could be achieved circumventing its suppressive effects on T cell proliferation (27), thus suggesting that BEA can be a very promising candidate of vaccine adjuvants and cancer immunotherapy. In addition, BEA has been reported to neutralize the ATP-binding cassette (ABC) transporters, which contributes to multi-drug resistance in human, nematodes and arthropods (15, 45). Therefore, combinational therapy using BEA and other drugs can overcome multidrug resistance.

In summary, our data revealed a novel function of BEA on DCs in activating inflammatory cytokine and chemokine production via activating the TLR4 signaling pathway. Our findings suggest BEA can be exploited in the field of vaccine adjuvants and cancer immunotherapy.

## Author contributions

S.S. conceived and supervised the study. X.Y. performed the experiments, analyzed the results, X.Y., S.S., J.Q and M.U. wrote the manuscript. S.A. performed cell sorting and gave suggestions to the project. V.S. performed qRT-PCR experiments. L.R. screened natural products. C.K., H.W., P.L., U.K., J.S. and H.I. provided mice and isolated bone marrow. M.F. and R.K. isolated BEA and performed purity analyses. C.A. performed the Multiplex immunoassays. T.W and K.K performed RNA-seq experiments. X.Y., M.Z., J.D., S.B. and A.H. analyzed the RNA-seq data.

## Data availability statement

RNA sequencing data in this study have been deposited in NCBI Gene Expression Omnibus under the accession number GSE192689. Related website is: https://www.ncbi.nlm.nih.gov/geo/query/acc.cgi?acc=GSE192689.

## Acknowledgement

Computational support of the ‘Zentrum für Informations-und Medientechnologie’, especially the HPC team (High Performance Computing) at the Heinrich-Heine University is acknowledged. This work was funded by the Deutsche Forschungsgemeinschaft (DFG, German Research Foundation) DFG-270650915/GRK 2158 to S.S., by the Deutsche Forschungsgemeinschaft (DFG; German Research Foundation) -398066876/GRK 2485/1 to U.K., by the Deutsche Forschungsgemeinschaft (DFG; German Research Foundation) DFG-158989968 – DFB 900-B2 to U.K., by the Deutsche Forschungsgemeinschaft (DFG; German Research Foundation) under Germany’s Excellence Strategy – EXC 2155 “RESIST” – Project ID 39087428 to U.K.

---

**supplementary Figure 1.**
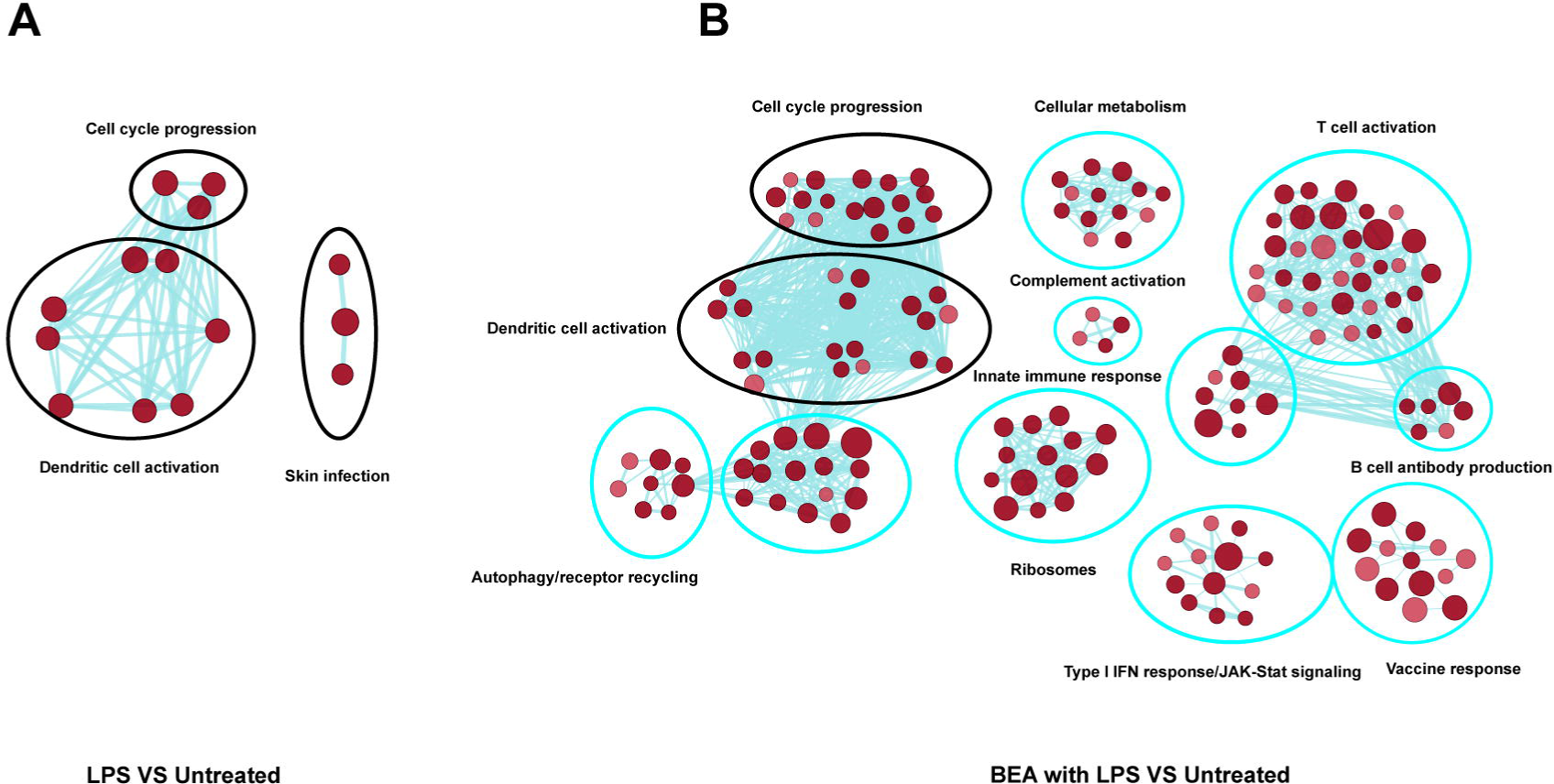
**(A)** Cytoscape representation of significantly enriched signatures in LPS treated BMDCs compared with untreated BMDCs. **(B)** Cytoscape representation of significantly enriched signatures in BEA with LPS treated BMDCs compared with untreated BMDCs.

## References

1. Banchereau J, Briere F, Caux C, Davoust J, Lebecque S, Liu YJ, et al. Immunobiology of dendritic cells. Annu Rev Immunol. 2000;18:767–811.

2. Liu J, Zhang X, Cheng Y, Cao X. Dendritic cell migration in inflammation and immunity. Cell Mol Immunol. 2021;18(11):2461–71.

3. Steinman RM. Decisions about dendritic cells: past, present, and future. Annu Rev Immunol. 2012;30:1–22.

4. Iwasaki A, Medzhitov R. Toll-like receptor control of the adaptive immune responses. Nat Immunol. 2004;5(10):987–95.

5. Behzadi P, Garcia-Perdomo HA, Karpinski TM. Toll-Like Receptors: General Molecular and Structural Biology. J Immunol Res. 2021;2021:9914854.

6. Piqueras B, Connolly J, Freitas H, Palucka AK, Banchereau J. Upon viral exposure, myeloid and plasmacytoid dendritic cells produce 3 waves of distinct chemokines to recruit immune effectors. Blood. 2006;107(7):2613–8.

7. Foti M, Granucci F, Aggujaro D, Liboi E, Luini W, Minardi S, et al. Upon dendritic cell (DC) activation chemokines and chemokine receptor expression are rapidly regulated for recruitment and maintenance of DC at the inflammatory site. Int Immunol. 1999;11(6):979–86.

8. Hilligan KL, Ronchese F. Antigen presentation by dendritic cells and their instruction of CD4+ T helper cell responses. Cell Mol Immunol. 2020;17(6):587–99.

9. Trinchieri G. Interleukin-12 and the regulation of innate resistance and adaptive immunity. Nat Rev Immunol. 2003;3(2):133–46.

10. Logrieco A, Moretti A, Castella G, Kostecki M, Golinski P, Ritieni A, et al. Beauvericin production by Fusarium species. Appl Environ Microbiol. 1998;64(8):3084–8.

11. Peczynska-Czoch W, Urbanczyk MJ, Balazy S. Formation of beauvericin by selected strains of Beauveria bassiana. Arch Immunol Ther Exp (Warsz). 1991;39(1-2):175–9.

12. Han X, Xu W, Zhang J, Xu J, Li F. Natural Occurrence of Beauvericin and Enniatins in Corn-and Wheat-Based Samples Harvested in 2017 Collected from Shandong Province, China. Toxins. 2018;11(1).

13. Juan C, Manes J, Raiola A, Ritieni A. Evaluation of beauvericin and enniatins in Italian cereal products and multicereal food by liquid chromatography coupled to triple quadrupole mass spectrometry. Food chemistry. 2013;140(4):755–62.

14. Wu Q, Patocka J, Nepovimova E, Kuca K. A Review on the Synthesis and Bioactivity Aspects of Beauvericin, a Fusarium Mycotoxin. Front Pharmacol. 2018;9:1338.

15. Al Khoury C, Nemer N, Nemer G. Beauvericin potentiates the activity of pesticides by neutralizing the ATP-binding cassette transporters in arthropods. Scientific reports. 2021;11(1):10865.

16. Xu L, Wang J, Zhao J, Li P, Shan T, Wang J, et al. Beauvericin from the endophytic fungus, Fusarium redolens, isolated from Dioscorea zingiberensis and its antibacterial activity. Nat Prod Commun. 2010;5(5):811–4.

17. Shin CG, An DG, Song HH, Lee C. Beauvericin and enniatins H, I and MK1688 are new potent inhibitors of human immunodeficiency virus type-1 integrase. J Antibiot (Tokyo). 2009;62(12):687–90.

18. Lim HN, Jang JP, Shin HJ, Jang JH, Ahn JS, Jung HJ. Cytotoxic Activities and Molecular Mechanisms of the Beauvericin and Beauvericin G1 Microbial Products against Melanoma Cells. Molecules. 2020;25(8).

19. Ferrer E, Juan-Garcia A, Font G, Ruiz MJ. Reactive oxygen species induced by beauvericin, patulin and zearalenone in CHO-K1 cells. Toxicol In Vitro. 2009;23(8):1504–9.

20. Yoo S, Kim MY, Cho JY. Beauvericin, a cyclic peptide, inhibits inflammatory responses in macrophages by inhibiting the NF-kappaB pathway. The Korean journal of physiology & pharmacology : official journal of the Korean Physiological Society and the Korean Society of Pharmacology. 2017;21(4):449–56.

21. Reinhardt RL, Hong S, Kang SJ, Wang ZE, Locksley RM. Visualization of IL-12/23p40 in vivo reveals immunostimulatory dendritic cell migrants that promote Th1 differentiation. J Immunol. 2006;177(3):1618–27.

22. Love MI, Huber W, Anders S. Moderated estimation of fold change and dispersion for RNA-seq data with DESeq2. Genome Biol. 2014;15(12):550.

23. Mootha VK, Lindgren CM, Eriksson KF, Subramanian A, Sihag S, Lehar J, et al. PGC-1alpha-responsive genes involved in oxidative phosphorylation are coordinately downregulated in human diabetes. Nat Genet. 2003;34(3):267–73.

24. Subramanian A, Tamayo P, Mootha VK, Mukherjee S, Ebert BL, Gillette MA, et al. Gene set enrichment analysis: a knowledge-based approach for interpreting genome-wide expression profiles. Proc Natl Acad Sci U S A. 2005;102(43):15545–50.

25. Shannon P, Markiel A, Ozier O, Baliga NS, Wang JT, Ramage D, et al. Cytoscape: a software environment for integrated models of biomolecular interaction networks. Genome Res. 2003;13(11):2498–504.

26. Merico D, Isserlin R, Stueker O, Emili A, Bader GD. Enrichment map: a network-based method for gene-set enrichment visualization and interpretation. PLoS One. 2010;5(11):e13984.

27. Wu XF, Xu R, Ouyang ZJ, Qian C, Shen Y, Wu XD, et al. Beauvericin ameliorates experimental colitis by inhibiting activated T cells via downregulation of the PI3K/Akt signaling pathway. PloS one. 2013;8(12):e83013.

28. Deng SL, Zhang BL, Reiter RJ, Liu YX. Melatonin Ameliorates Inflammation and Oxidative Stress by Suppressing the p38MAPK Signaling Pathway in LPS-Induced Sheep Orchitis. Antioxidants (Basel). 2020;9(12).

29. Anand G, Perry AM, Cummings CL, Raymond E, Clemens RA, Steed AL. Surface Proteins of SARS-CoV-2 Drive Airway Epithelial Cells to Induce IFN-Dependent Inflammation. J Immunol. 2021.

30. Krummen M, Balkow S, Shen L, Heinz S, Loquai C, Probst HC, et al. Release of IL-12 by dendritic cells activated by TLR ligation is dependent on MyD88 signaling, whereas TRIF signaling is indispensable for TLR synergy. Journal of leukocyte biology. 2010;88(1):189–99.

31. Shen H, Tesar BM, Walker WE, Goldstein DR. Dual signaling of MyD88 and TRIF is critical for maximal TLR4-induced dendritic cell maturation. Journal of immunology. 2008;181(3):1849–58.

32. Bandow K, Kusuyama J, Shamoto M, Kakimoto K, Ohnishi T, Matsuguchi T. LPS-induced chemokine expression in both MyD88-dependent and -independent manners is regulated by Cot/Tpl2-ERK axis in macrophages. FEBS letters. 2012;586(10):1540–6.

33. Hemmi H, Akira S. TLR signalling and the function of dendritic cells. Chem Immunol Allergy. 2005;86:120–35.

34. Kawai T, Akira S. Toll-like receptor downstream signaling. Arthritis Res Ther. 2005;7(1):12–9.

35. Netea MG, Warris A, Van der Meer JW, Fenton MJ, Verver-Janssen TJ, Jacobs LE, et al. Aspergillus fumigatus evades immune recognition during germination through loss of toll-like receptor-4-mediated signal transduction. The Journal of infectious diseases. 2003;188(2):320–6.

36. Ficheux AS, Sibiril Y, Parent-Massin D. Effects of beauvericin, enniatin b and moniliformin on human dendritic cells and macrophages: an in vitro study. Toxicon. 2013;71:1–10.

37. Vogl T, Tenbrock K, Ludwig S, Leukert N, Ehrhardt C, van Zoelen MA, et al. Mrp8 and Mrp14 are endogenous activators of Toll-like receptor 4, promoting lethal, endotoxin-induced shock. Nat Med. 2007;13(9):1042–9.

38. Tsan MF, Gao B. Endogenous ligands of Toll-like receptors. Journal of leukocyte biology. 2004;76(3):514–9.

39. Al-Ofi EA, Al-Ghamdi BS. High-mobility group box 1, an endogenous ligand of toll-like receptors 2 and 4, induces astroglial inflammation via nuclear factor kappa B pathway. Folia Morphol (Warsz). 2019;78(1):10–6.

40. Kumar S, Sunagar R, Gosselin E. Bacterial Protein Toll-Like-Receptor Agonists: A Novel Perspective on Vaccine Adjuvants. Front Immunol. 2019;10:1144.

41. Maisonneuve C, Bertholet S, Philpott DJ, De Gregorio E. Unleashing the potential of NOD-and Toll-like agonists as vaccine adjuvants. Proc Natl Acad Sci U S A. 2014;111(34):12294–9.

42. Taleghani N, Bozorg A, Azimi A, Zamani H. Immunogenicity of HPV and HBV vaccines: adjuvanticity of synthetic analogs of monophosphoryl lipid A combined with aluminum hydroxide. APMIS. 2019;127(3):150–7.

43. Shetab Boushehri MA, Lamprecht A. TLR4-Based Immunotherapeutics in Cancer: A Review of the Achievements and Shortcomings. Mol Pharm. 2018;15(11):4777–800.

44. Cluff CW. Monophosphoryl lipid A (MPL) as an adjuvant for anti-cancer vaccines: clinical results. Adv Exp Med Biol. 2010;667:111–23.

45. Wu C, Chakrabarty S, Jin M, Liu K, Xiao Y. Insect ATP-Binding Cassette (ABC) Transporters: Roles in Xenobiotic Detoxification and Bt Insecticidal Activity. International journal of molecular sciences. 2019;20(11).

